# Transient patterns of functional dysconnectivity in youth with psychosis spectrum symptoms

**DOI:** 10.1101/426932

**Authors:** Eva Mennigen, Dietsje D. Jolles, Catherine E. Hegarty, Mohan Gupta, Maria Jalbrzikowski, Loes M. Olde Loohuis, Roel A. Ophoff, Katherine H. Karlsgodt, Carrie E. Bearden

## Abstract

Psychosis spectrum disorders are conceptualized as neurodevelopmental disorders accompanied by disruption of large-scale functional brain networks. Both static and dynamic dysconnectivity have been described in patients with schizophrenia and, more recently, in help-seeking individuals at clinical high-risk for psychosis. Less is known, however, about developmental aspects of dynamic functional network connectivity (FNC) associated with psychotic symptoms (PS) in the general population. Here, we investigate resting state fMRI data using established dynamic FNC methods in the Philadelphia Neurodevelopmental Cohort (ages 8-22), including 129 participants experiencing PS and 452 participants without PS (non-PS).

Applying a sliding window approach and k-means clustering, 5 dynamic states with distinct whole-brain connectivity patterns were identified. PS-associated dysconnectivity was most prominent in states characterized by synchronization or antagonism of the default mode network (DMN) and cognitive control (CC) domains. Hyperconnectivity between DMN, salience, and CC domains in PS youth only occurred in a state characterized by synchronization of the DMN and CC domains, a state that also becomes less frequent with age. However, dysconnectivity of the sensorimotor and visual systems in PS youth was revealed in other transient states completing the picture of whole-brain dysconnectivity patterns associated with PS.

Overall, state-dependent dysconnectivity was observed in PS youth, providing the first evidence that disruptions of dynamic functional connectivity are present across a broader psychosis continuum.

## Introduction

Substantial evidence now indicates that psychotic symptoms (PS) occur on a continuum ranging from sub-threshold PS to full-blown psychotic disorders such as schizophrenia.^1–3^ Clinically, PS include abnormalities of perception, emotion, and cognition that vary in severity, frequency, and level of conviction across the psychosis spectrum.^4,5^ Traditionally, individuals on the severe end of this continuum have been studied. But more recently, there has been increasing interest in individuals experiencing a broader spectrum of PS. First, because they are at increased risk of progressing to overt illness,^6,7^ but secondly because they offer the opportunity to explore neural changes in the absence of confounds from medication or disease chronicity.

The psychosis continuum is considered to have neurodevelopmental underpinnings concomitant with altered brain and cognitive maturation.^8–12^ Symptoms of many psychiatric illnesses typically appear during adolescence, a sensitive period of brain development,^13–15^ and frequency of PS peaks in late childhood and adolescence.^2,16^ Therefore, brain imaging studies of youth experiencing PS are likely to be informative regarding neural substrates of developmental vulnerability to psychosis. Publicly available data from the Philadelphia Neurodevelopmental Cohort (PNC) utilized in the current study offer an unprecedented opportunity to study neural substrates of PS from late-childhood through adolescence and early adulthood, overlapping with critical periods for the onset of many neuropsychiatric disorders.^17,18^

There is now a wealth of evidence that disruption of large-scale synchronized neural connectivity plays a role in the pathophysiology of schizophrenia.^19–21^ Functional connectivity describes the correlated temporal fluctuations of distant brain areas, and is often assessed during resting state functional magnetic resonance imaging (rs-fMRI) while participants are not engaged in a particular task.^22–24^ In terms of static functional connectivity, which reflects the average connectivity across the entire resting state scan, previous findings in PS youth in this cohort include hyperconnectivity within the default mode network (DMN) that was associated with poorer cognitive performance, and hypoconnectivity within the cognitive control (CC) domain.^11^ These patterns resemble those observed in patients with overt schizophrenia, as well as in help-seeking individuals at clinical high-risk (CHR) for psychosis.

Only recently has it emerged that functional connectivity is not static, but rather a dynamic process that exhibits considerable fluctuations across the duration of a typical resting state scan.^25–27^ Indeed, dynamic or state effects are as important as static or trait effects in determining individual functional connectivity patterns. Greater variability in network activity is associated with increased capacity for information processing,^28^ and thus may index better overall ‘brain health’.^29,30^ With the emergence of new methods, we are now poised to explore the dynamics of functional dysconnectivity related to PS.^26,31–34^

Recently, we investigated dynamic functional network connectivity (FNC) utilizing a sliding window approach^31^ to identify recurring whole brain connectivity patterns in treatment-seeking CHR youth.^35^ Abnormalities were only observed within specific dynamic states, and overall fluctuations of connectivity across dynamic states in CHR individuals were reduced relative to healthy controls. Further, CHR individuals exhibit qualitatively similar, but milder, dysconnectivity relative to patients with schizophrenia.^36^ Applying a different approach to capture dynamic aspects of functional connectivity,^33^ Barber et al. investigated individuals endorsing PS who were otherwise healthy;^37^ individuals with PS spent more time in states that showed intra-DMN hypoconnectivity, consistent with findings in patients with overt schizophrenia.^38,39^

The aim of the current study was to investigate whole-brain dynamic FNC and associated summary metrics in PS youth relative to their peers who do not experience PS (non-PS) across late childhood, adolescence, and early adulthood.

## Methods

### Study participants

An ethnically and socioeconomically diverse community sample of participants aged 8 to 22 years was included in the PNC study. This study was not designed to ascertain individuals with particular neuropsychiatric disorders, but instead recruited participants broadly from the Children’s Hospital of Philadelphia.

Study participants (n=9,428) completed a computerized structured interview (GOASSESS) that included a psychopathology screening based on the National Institute of Mental Health Genetic Epidemiology Research Branch Kiddie – Schedule for Affective Disorders and Schizophrenia (K-SADS)^40^ and a computerized neurocognitive battery (CNB)^41^. Multimodal MRI was acquired for a subsample of participants (n=1,445).^42^

Out of 799 participants with rs-fMRI scans, imaging data of 581 participants with available age and sex information passed quality control. Demographics for this sample are summarized in Table 1.

### Clinical Interview

GOASSESS was developed to allow for a large ‘throughput’ of participants. It provides screen-level symptom and episode information^43^ and is based on the K-SADS. See Supplementary Material for additional information.

### Psychosis Spectrum Classification

We identified PS individuals in the cohort according to criteria used by Calkins et al.^44^ Briefly, PS were rated based on the PRIME Screen-Revised^45^ assessing positive symptoms, the K-SADS^46^ for hallucinations and delusional symptoms, and the Scale of Prodromal Syndromes^47^ assessing negative and disorganized symptoms (see Supplementary Material).

### Resting state fMRI data and preprocessing

Eyes-open rs-fMRI data were collected on a single scanner with 3T field strength over 6.2 minutes. FMRIB Software Library (FSL; https://fsl.fmrib.ox.ac.uk/fsl) and Analysis of Functional NeuroImages (AFNI; https://afni.nimh.nih.gov) tools were used for functional preprocessing that included slice time correction, motion correction, grand mean scaling, and smoothing (6mm kernel). Table 1 contains a comparison of motion parameters between groups.

See Supplementary Material for additional information.

**Table 1:** Demographics and motion parameters

| Group | Age<br>(SD) | Sex<br>(%<br>female) | Ethnicity in % |  |  | Maternal<br>education | Relative<br>movement | Maximum<br>movement | Spike<br>count |
| --- | --- | --- | --- | --- | --- | --- | --- | --- | --- |
|  |  |  | AA | EA | Other |  |  |  |  |
| **Non-**<br>**PS**<br>**n=452** | 15.2<br>(3.2) | 55.3 | 35.8<sup>+</sup> | 54.2<sup>+</sup> | 10.0<sup>+</sup> | 14.4 (2.5) * | 0.59 (0.29)<br>* | 0.7 (0.59) | 4.38<br>(5.3) |
| **PS**<br>**n=129** | 15.00<br>(2.8) | 56.6 | 57.4<sup>+</sup> | 31.0<sup>+</sup> | 11.6<sup>+</sup> | 13.8 (2.2) * | 0.65 (0.29)<br>* | 0.77 (0.57) | 5.24<br>(5.6) |

Significant group difference ( $p < 0.05$ ): \* two-sample t-test, <sup>+</sup> Chi-square test; PS – psychosis spectrum; AA – African-American, EA – European-American

### Group Independent Component Analysis

Independent component analysis (ICA) is a special case of blind source separation and is widely applied to imaging data. RS-fMRI data were decomposed into cortical and subcortical components using a high model order group-level spatial ICA^48^ using the Group ICA fMRI toolbox (GIFT, http://mialab.mrn.org/software/gift).

Two independent raters (EM, DDJ) evaluated 59 out of 100 components as intrinsic connectivity networks (ICNs) based on the following criteria:^49^ ICNs show peak activation in gray matter with no or minimal overlap with white matter, ventricles, or non-brain structures and ICNs show maximal power in lower frequencies (< 0.1 Hz). ICNs were assigned to the following 9 functional domains based on their anatomical location and prior scientific literature utilizing the automated anatomic labeling atlas^50^ and neurosynth.org: subcortical, salience, auditory, sensorimotor, visual, cognitive control (CC), DMN, limbic, and cerebellum. Figure 1 shows the 9 functional domains with their assigned ICNs. See Supplementary Material for additional information.

**Figure 1:**
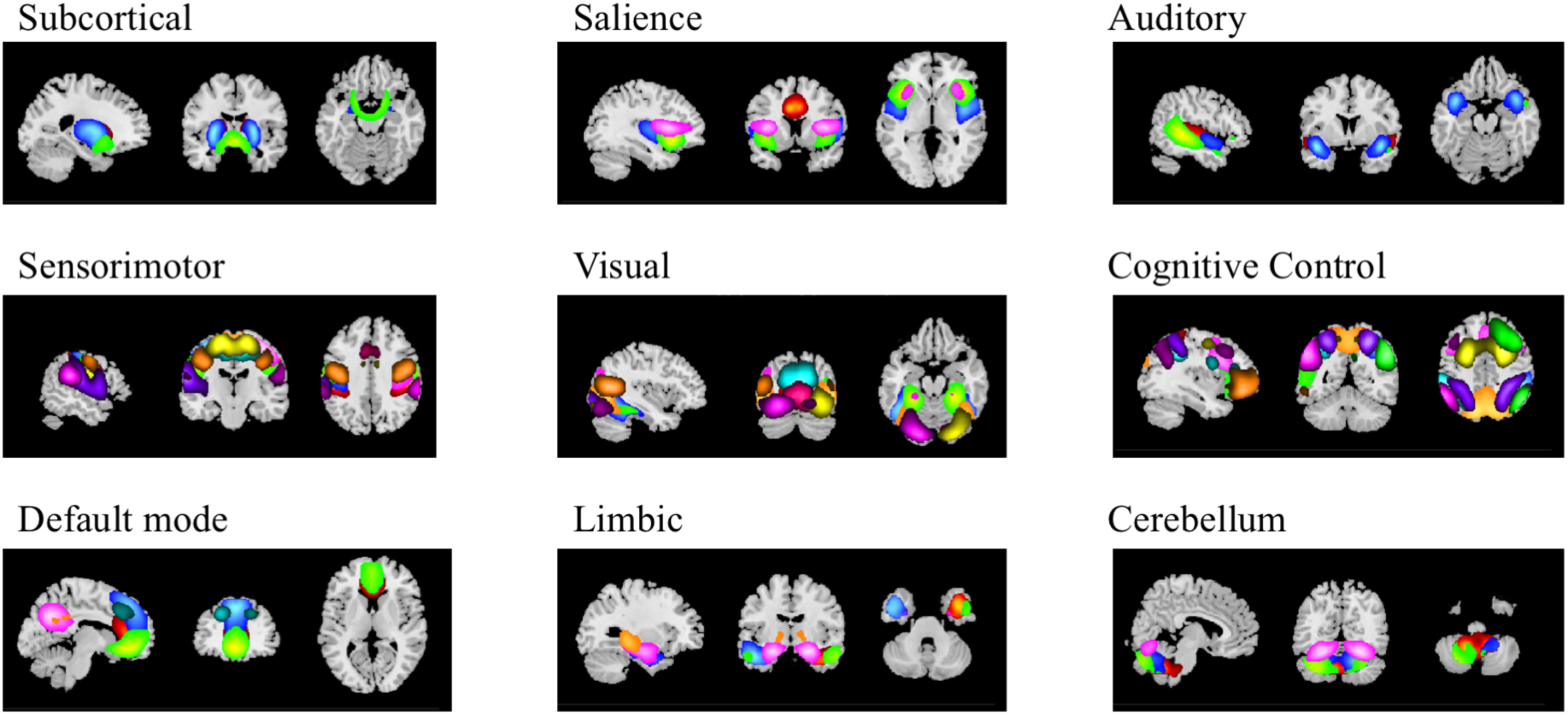
Nine functional domains and their assigned intrinsic connectivity networks

### Dynamic FNC

We applied a sliding temporal window approach to capture changes of whole-brain connectivity (Figure S1).^31^ Briefly, a tapered window slides across concatenated time courses and for each window an FNC matrix consisting of ICN-to-ICN Pearson’s correlations was calculated.

Next, from each participant, windows with the highest variance in FNC (‘local extrema’) were chosen to initialize clustering. K-means clustering was first performed on the local extrema with varying numbers of clusters *k* (2-20): The ratio of within- to between-cluster distances was plotted for each *k*. The turning point in the graph where the amount of additionally explained variance becomes marginal, and therefore reflecting the optimal number of clusters (‘elbow criterion’), was five, which is typical for this type of analysis.^51^

These five cluster centroids were then used as starting points to cluster all windowed FNC matrices in such a way that each windowed FNC matrix was assigned to the one cluster with which it was most highly correlated. For each participant, each dynamic state is represented by the element-wise median connectivity across all windows assigned to this particular state. The five dynamic states are shown in Figure 2.

**Figure 2:**
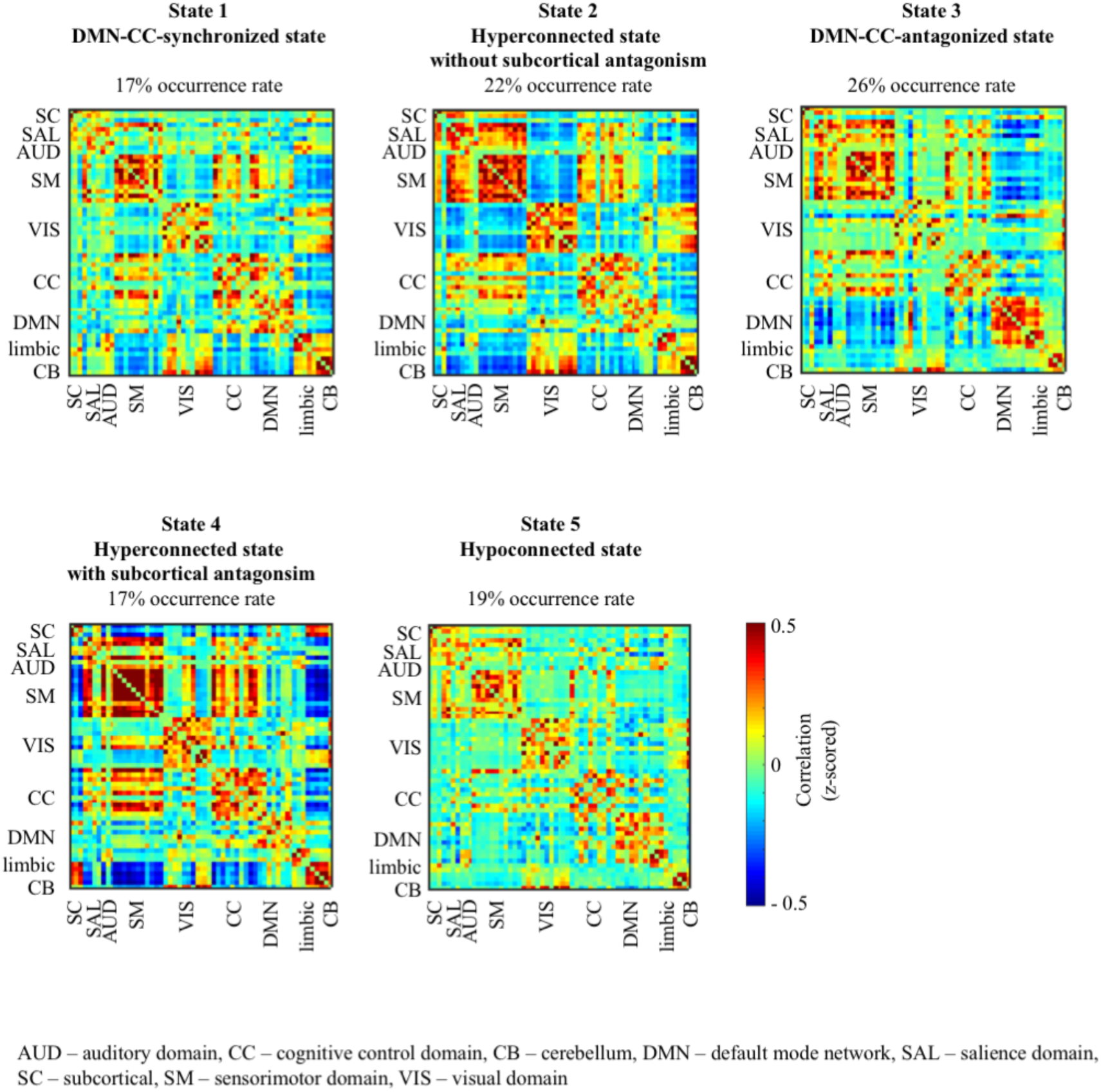
The five dynamic states identified, including their occurrence rates across all participants.

## Model selection

In order to account for important covariates but also to prevent the model from overfitting, we applied a multivariate backward model selection approach adapted from the MANCOVAN toolbox implemented in GIFT using the mSTEPWISE function.^49^ Assuming that each dynamic state may be influenced differently by the covariates, statistical models were generated for each state separately. The initial full model for all dynamic states included the following variables: group (non-PS vs. PS), sex, age, maternal education, and their interactions (see Supplementary Material).

The following models were selected for the five dynamic states:

- state 1: FNC ~ (group, sex, age, group * sex) * β + ε
- state 2: FNC ~ (sex, age, maternal education) * β + ε
- state 3: FNC ~ (group, sex, age, maternal education) * β + ε
- state 4: FNC ~ (group, sex, age, maternal education, group * age, sex * maternal education) * β + ε
- state 5: FNC ~ (group, sex, age, maternal education, group * maternal education) * β + ε

The reduced models were then used for further univariate tests.^49^ Results were corrected for a false discovery rate (FDR) at q = 0.05.

### Dynamic indices

Based on the distinction of five discrete states, summary metrics reflecting the dynamic behavior of FNC across the scan can be derived. The mean dwell time (MDT) reflects the average time an individual lingers in one particular state before switching to a different state; the fraction of time (FT) summarizes the time across the entire scan that an individual spends in one particular state.

We applied the same backward model selection procedure as for the dynamic FNC analysis with the same set of covariates. The reduced models for FT and MDT included the covariates sex, age, and maternal education but not group. Results were FDR corrected at q = 0.05.

## Results

### Dynamic FNC

The five dynamic states are shown in Figure 2; their distinct connectivity patterns are described individually below. With regard to connectivity differences, we focus on results for our primary variable of interest, group (PS vs. non-PS), which was included as factor in the reduced models of states 1,3,4, and 5. Figure 3 shows significant group effects across dynamic states.

**Figure 3:**
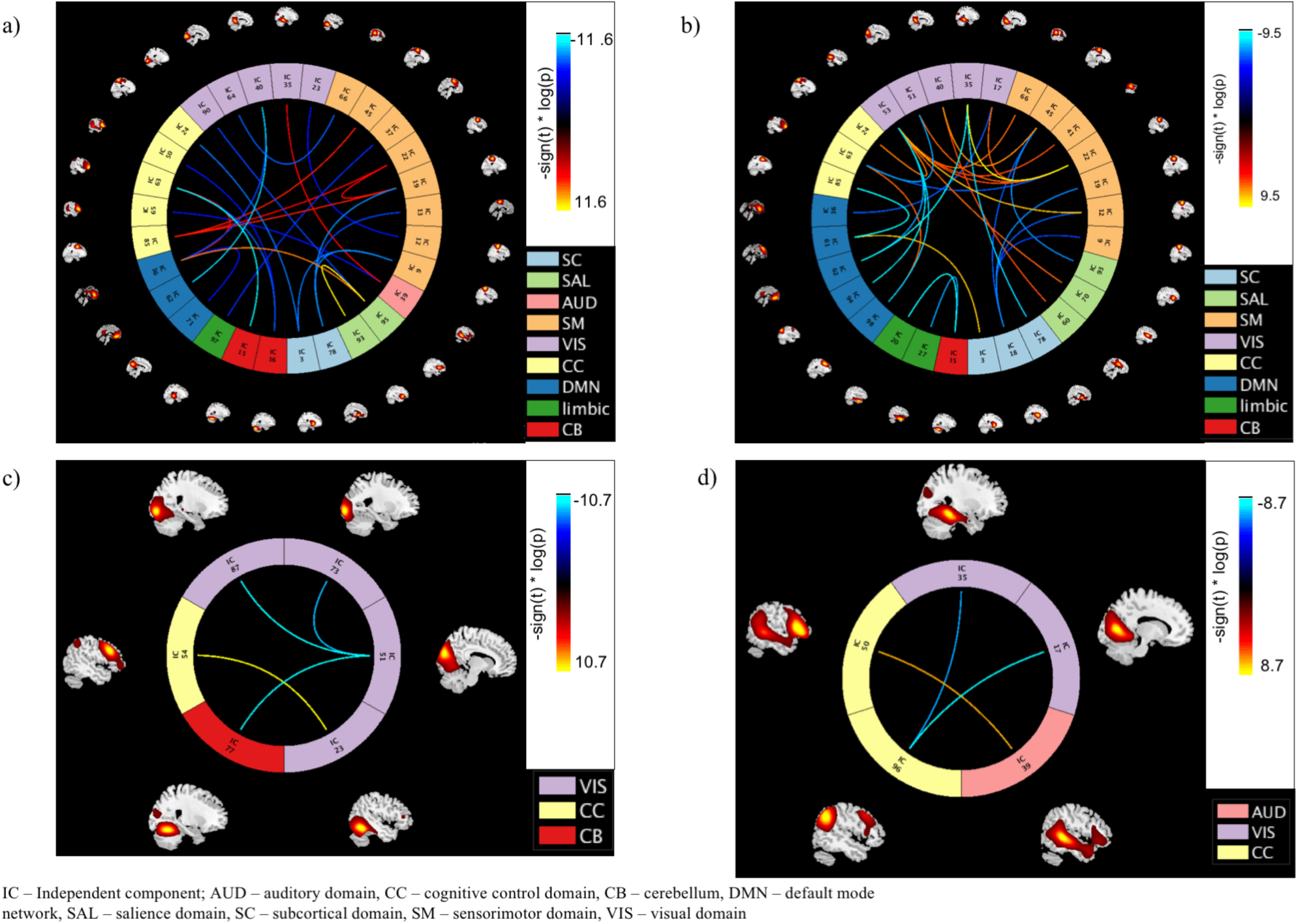
*ICN-to-ICN connections showing significant group effects in a) state 1, b) state 3, and c) state 5; d) ICN-to-ICN connections of significant group by age interaction effects in state 4. The scaling, -sign(t)* * *log(p), provides information on the effect size and direction. The cool color scale represents negative values, indicating hypoconnectivity (decreased positive correlation, or greater anti-correlation) in PS relative to non-PS youth; the hot color scale represents positive values indicating hyperconnectivity (increased positive correlation or less anti-correlation) in PS relative to non-PS youth*.

## State 1 DMN-CC domain-synchronized state

Across all participants, 17% of all windows were assigned to this state. For the most part, DMN and CC domains appear synchronized in this state: they show high positive connectivity with each other, and form one functional domain. Together they exhibit negative connectivity with the limbic domain and the cerebellum. Further, state 1 shows anti-correlation between the sensorimotor domain and limbic and cerebellar domains.

In this state, twenty-four ICN-to-ICN connectivity pairs show a significant group effect (Figure 3a, Table 2). In general, PS youth exhibit reduced connectivity between the CC domain with multiple other domains (auditory, cerebellar, subcortical) as well as between the sensorimotor domain with visual and subcortical domains and reduced intra-domain connectivity within the DMN. In contrast, increased inter-domain connectivity in PS relative to non-PS youth is observed between the CC domain with the DMN and the sensorimotor domains, as well as between auditory and visual domains, salience domain and DMN, and increased *intra*-domain connectivity within the salience domain.

**Table 2:**
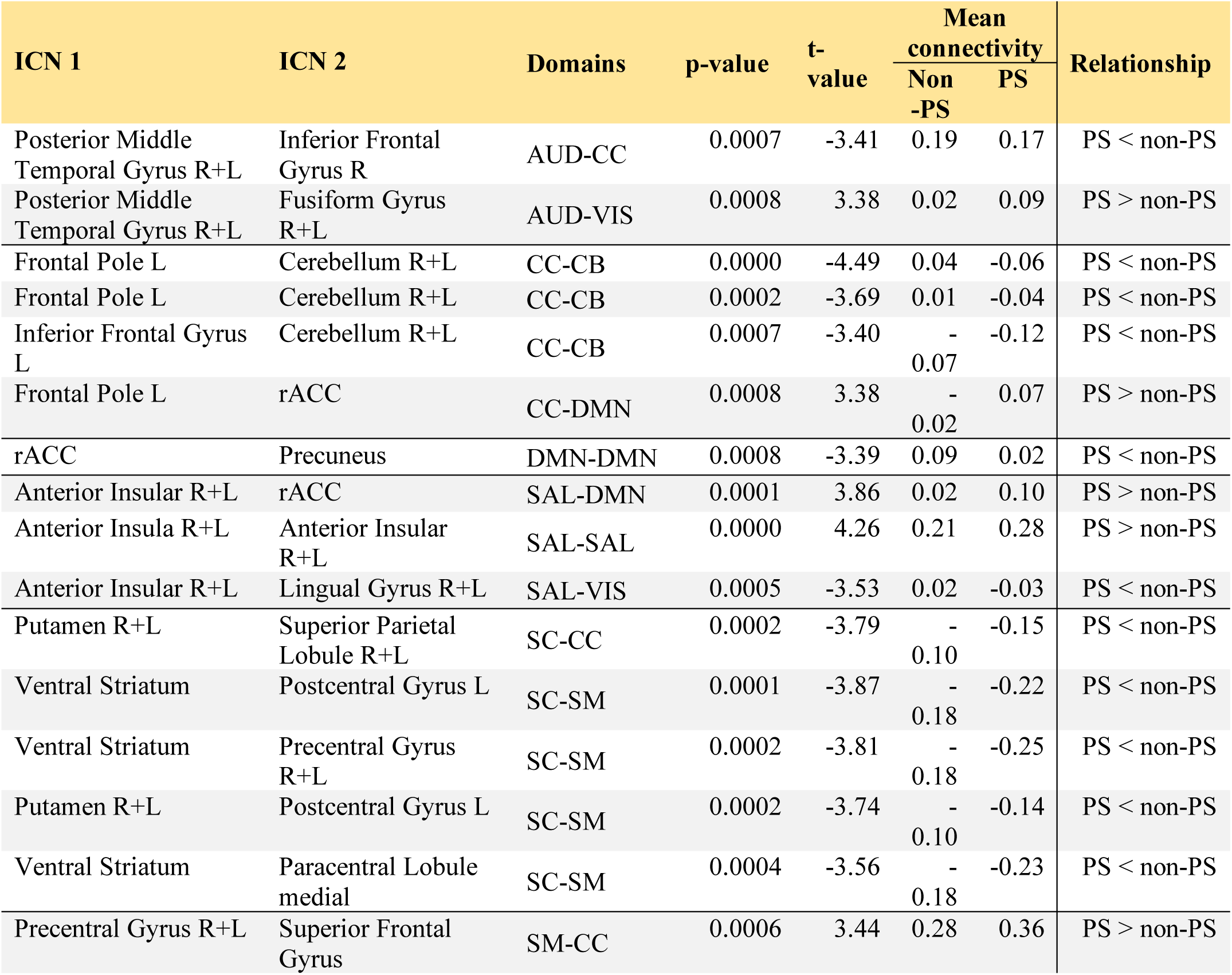
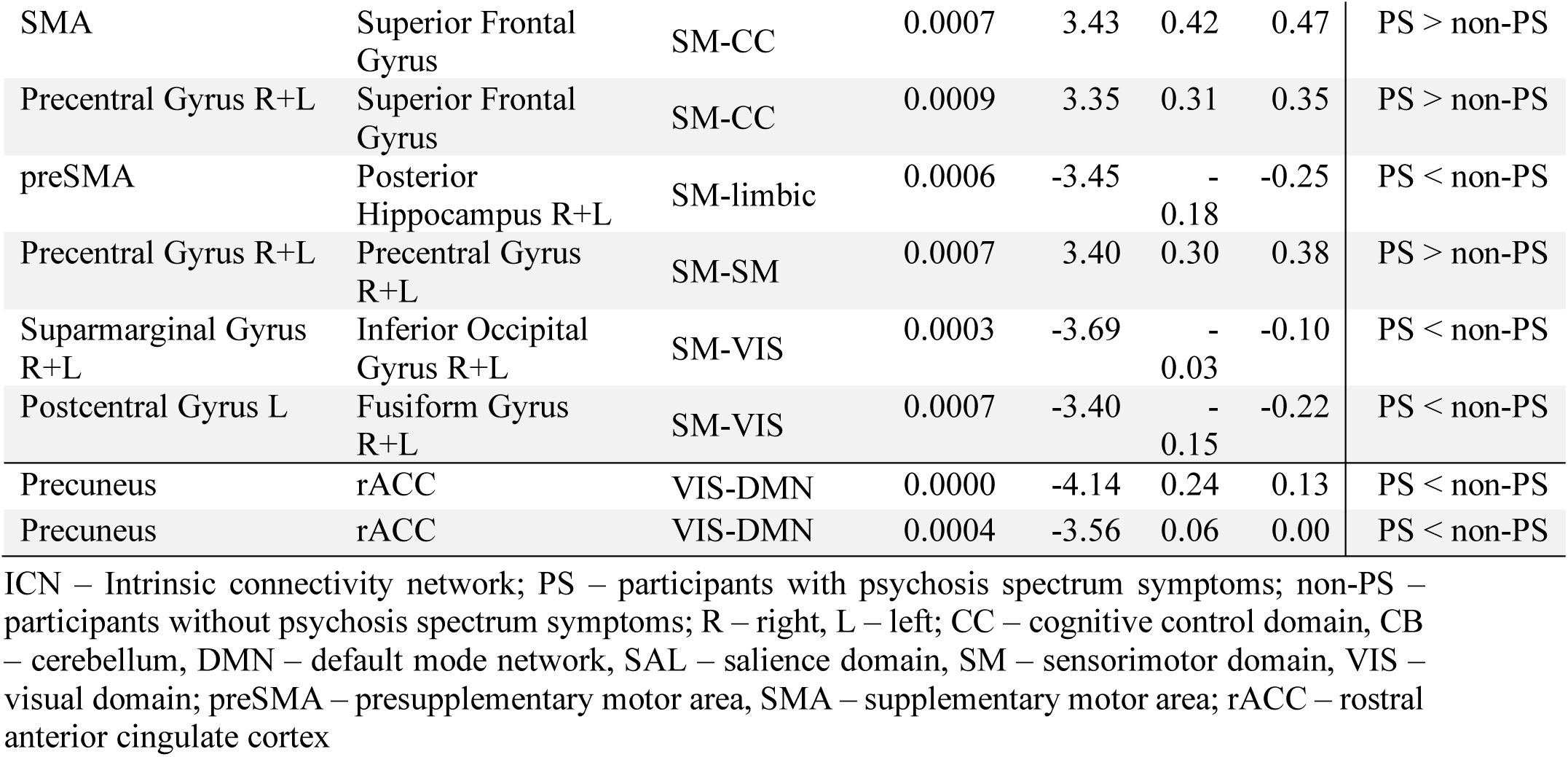
ICN-to-ICN connectivity pairs that show significant group effects in state 1 (DMN-CC-synchronized state), ordered by domains.

## State 2 Hyperconnected state without subcortical antagonism

State 2 is characterized by increased intra-domain connectivity, particularly in the salience, sensorimotor, and cerebellar domains. The visual domain is strongly anticorrelated with the sensorimotor, salience, and subcortical domains. CC and DMN domains appear synchronized. 22% of all windowed FNC matrices were assigned to this state.

## State 3 DMN-CC domain-antagonized state

In this state, each functional domain shows positive intra-domain connectivity with the exception of the visual domain. The DMN exhibits strong anti-correlation with the CC, salience, and sensorimotor domains. 26% of windowed FNC matrices were clustered into this pattern.

In state 3, 28 ICN-to-ICN connectivity pairs exhibit significant differences between groups (Figure 3b, Table 3). Here, connectivity involving the sensorimotor and visual domains is particularly affected in PS youth. PS participants show decreased inter-domain connectivity relative to non-PS youth of the sensorimotor domain with salience and subcortical domains, as well as between the visual domain and the DMN, and between the limbic and cerebellar domains. However, PS youth show relatively increased connectivity between the visual domain with sensorimotor and salience domains, and between the DMN and subcortical domains.

**Table 3:** ICN-to-ICN connectivity pairs that show significant group effects in state 3 (DMN-CC-antagonized state), ordered by domains.

| ICN 1 | ICN 2 | Domains | p-value | t-value | Mean connectivity |  | Relationship |
| --- | --- | --- | --- | --- | --- | --- | --- |
|  |  |  |  |  | Non-PS | PS |  |
| Superior Frontal Gyrus | Superior Frontal Gyrus medial | CC-DMN | 0.0002 | -3.78 | -0.13 | -0.22 | PS < non-PS |
| Frontal Pole L | Middle Frontal Gyrus R+L | CC-DMN | 0.001 | 3.32 | 0.03 | 0.11 | PS > non-PS |
| Temporal Pole | Cerebellum R+L | Limbic-CB | 0.0001 | -4.00 | 0.04 | -0.04 | PS < non-PS |
| Temporal Pole | Cerebellum R+L | Limbic-CB | 0.0005 | -3.49 | 0.04 | -0.03 | PS < non-PS |
| Anterior Insular R+L | SMA | SAL-SM | 0.0007 | -3.43 | 0.07 | -0.01 | PS < non-PS |
| Insular Cortex R+L | Posterior Middle Temporal Gyrus R+L | SAL-VIS | 0.0011 | 3.3 | -0.05 | 0.05 | PS > non-PS |
| dACC | Precuneus | SAL-VIS | 0.0013 | 3.24 | -0.26 | -0.20 | PS > non-PS |
| Putamen R+L | Superior Frontal Gyrus medial | SC-DMN | 0.0002 | 3.69 | -0.09 | -0.01 | PS > non-PS |
| Putamen R+L | Precentral Gyrus R+L | SC-SM | 0.0008 | -3.23 | 0.12 | 0.06 | PS < non-PS |
| Putamen R+L | SMA | SC-SM | 0.0013 | -3.23 | 0.2 | 0.13 | PS < non-PS |
| Putamen R+L | Postcentral Gyrus L | SC-SM | 0.0013 | -3.23 | 0.13 | 0.07 | PS < non-PS |
| Ventral Striatum | Fusiform Gyrus R+L | SC-VIS | 0.0003 | -3.67 | 0.07 | -0.01 | PS < non-PS |
| Precentral Gyrus R+L | Frontal Pole L | SM-CC | 0.0013 | -3.24 | -0.01 | -0.08 | PS < non-PS |
| Precentral Gyrus R+L | Fusiform Gyrus R+L | SM-VIS | 0.0001 | 3.97 | -0.13 | -0.05 | PS > non-PS |
| Postcentral Gyrus L | Posterior Middle Temporal Gyrus R+L | SM-VIS | 0.0003 | 3.62 | -0.01 | 0.09 | PS > non-PS |
| SMA | Precuneus | SM-VIS | 0.0005 | 3.5 | -0.29 | -0.22 | PS > non-PS |
| Supramarginal Gyrus R+L | Posterior Middle Temporal Gyrus R+L | SM-VIS | 0.0009 | 3.35 | 0.04 | 0.15 | PS > non-PS |
| Precentral Gyrus R+L | Cuneus | SM-VIS | 0.001 | 3.31 | 0.06 | 0.13 | PS > non-PS |
| Precentral Gyrus R+L | Lingual Gyrus R+L | SM-VIS | 0.001 | 3.3 | -0.03 | 0.05 | PS > non-PS |
| Precentral Gyrus R+L | Posterior Middle Temporal Gyrus R+L | SM-VIS | 0.0012 | 3.26 | -0.03 | 0.06 | PS > non-PS |
| Supramarginal Gyrus L | Posterior Middle Temporal Gyrus R+L | SM-VIS | 0.0014 | 3.22 | -0.01 | 0.09 | PS > non-PS |
| Cuneus | Frontal Pole L | VIS-CC | 0.0005 | -3.49 | -0.07 | -0.15 | PS < non-PS |
| Posterior Middle Temporal Gyrus R+L | Superior Parietal Lobule R+L | VIS-CC | 0.0006 | 3.45 | 0.03 | 0.13 | PS > non-PS |
| Fusiform Gyrus R+L | rACC | VIS-DMN | 0.0002 | -3.81 | 0.08 | -0.01 | PS < non-PS |
| Posterior Middle Temporal Gyrus R+L | Angular Gyrus R+L | VIS-DMN | 0.0002 | -3.72 | 0.09 | -0.02 | PS < non-PS |
| Posterior Middle Temporal Gyrus R+L | Middle Frontal Gyrus R+L | VIS-DMN | 0.0003 | -3.69 | 0.03 | -0.07 | PS < non-PS |
| Posterior Middle Temporal Gyrus R+L | rACC | VIS-DMN | 0.0006 | -3.48 | 0.07 | -0.04 | PS < non-PS |
| Lingual Gyrus R+L | rACC | VIS-DMN | 0.0013 | -3.24 | 0.00 | -0.06 | PS < non-PS |

ICN – Intrinsic connectivity network; PS – participants with psychosis spectrum symptoms; non-PS – participants without psychosis spectrum symptoms; R – right, L – left; AUD – auditory domain, CC – cognitive control domain, CB – cerebellum, DMN – default mode network, SAL – salience domain, SM – sensorimotor domain, VIS – visual domain; rACC – rostral anterior cingulate cortex, dACC – dorsal anterior cingulate cortex, SMA – supplementary motor area

## State 4 Hyperconnected state with subcortical antagonism

State 4 is characterized by increased intra-domain connectivity, particularly within the sensorimotor domain. Strong negative correlation is observed between the subcortical domain and sensorimotor, CC, and DMN domains, whereas connectivity between subcortical areas and the cerebellum is increased relative to the other states. The overall occurrence rate of this state was 17%.

Two ICN-to-ICN connectivity pairs show a significant group effect (Table 4, Figure 4). Since both pairs also exhibit a significant group by age interaction effect, only the interaction effect is considered. Specifically, PS youth exhibit age-associated decreases in connectivity between the right angular gyrus (CC domain) with both the lingual and fusiform gyri (visual domain), whereas non-PS participants show no change in connectivity with age. In contrast, connectivity increases with age between the posterior middle temporal gyrus (auditory domain) and the left inferior frontal gyrus (CC domain) in PS but not in non-PS youth.

**Table 4:** ICN-to-ICN connectivity pairs in state 4 (hyperconnected state with subcortical antagonism) that show: a) significant group effects, and b) group by age interaction effects.

| ICN1 | ICN2 | Domains | p-value | t-value | Mean connectivity |  | Relationship |
| --- | --- | --- | --- | --- | --- | --- | --- |
|  |  |  |  |  | Non-PS | PS |  |
| Lingual Gyrus R+L | Angular Gyrus R | VIS-DMN | 0.0001 | 3.84 | -0.12 | -0.11 | PS > non-PS |
| Fusiform Gyrus R+L | Angular Gyrus R | VIS-DMN | 0.001 | 3.32 | -0.13 | -0.12 | PS > non-PS |

a)
| ICN1 | ICN2 | Domains | p-value | t-value |
| --- | --- | --- | --- | --- |
| Lingual Gyrus R+L | Angular Gyrus R | VIS-CC | 0.0002 | -3.8 |
| Fusiform Gyrus R+L | Angular Gyrus R | VIS-CC | 0.001 | -3.29 |
| Posterior Middle Temporal Gyrus R+L | Inferior Frontal Gyrus L | AUD-CC | 0.0008 | 3.39 |

ICN – Intrinsic connectivity network; PS – participants with psychosis spectrum symptoms; non-PS – participants without psychosis spectrum symptoms; R – right, L – left; CC – cognitive control domain, DMN – default mode network, VIS – visual domain

**Figure 4:**
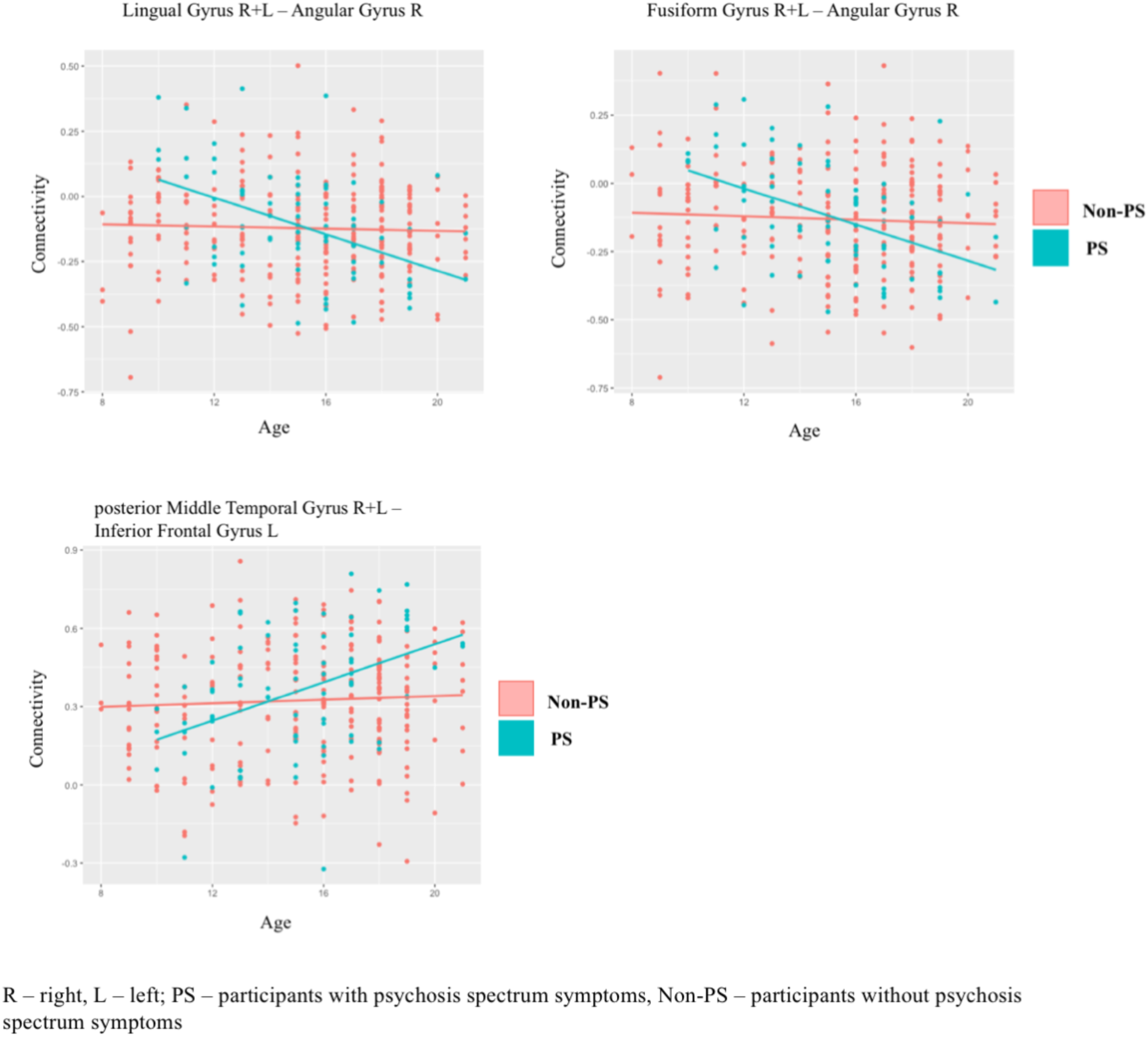
Scatterplots of the significant group by age interaction in state 4, showing decreased connectivity with age in PS relative to non-PS youth between two ICN-to-ICN pairs: lingual gyri – right angular gyrus and fusiform gyri – right angular gyrus. In contrast, PS youth show increased connectivity with age between posterior middle temporal gyri – left inferior frontal gyrus, relative to non-PS youth.

## State 5 Globally hypoconnected state

In this state, connectivity across domains appears diminished and functional domains are less distinguishable based on their intra-domain connectivity. Of all windowed FNC matrices, 19% were assigned to this state.

Four ICN-to-ICN connectivity pairs show significant differences between non-PS and PS groups (Table 5): connectivity within the visual domain is reduced in PS, whereas connectivity between fusiform gyri (visual domain) and middle frontal gyri (CC domain) is increased in PS relative to non-PS youth.

**Table 5:**
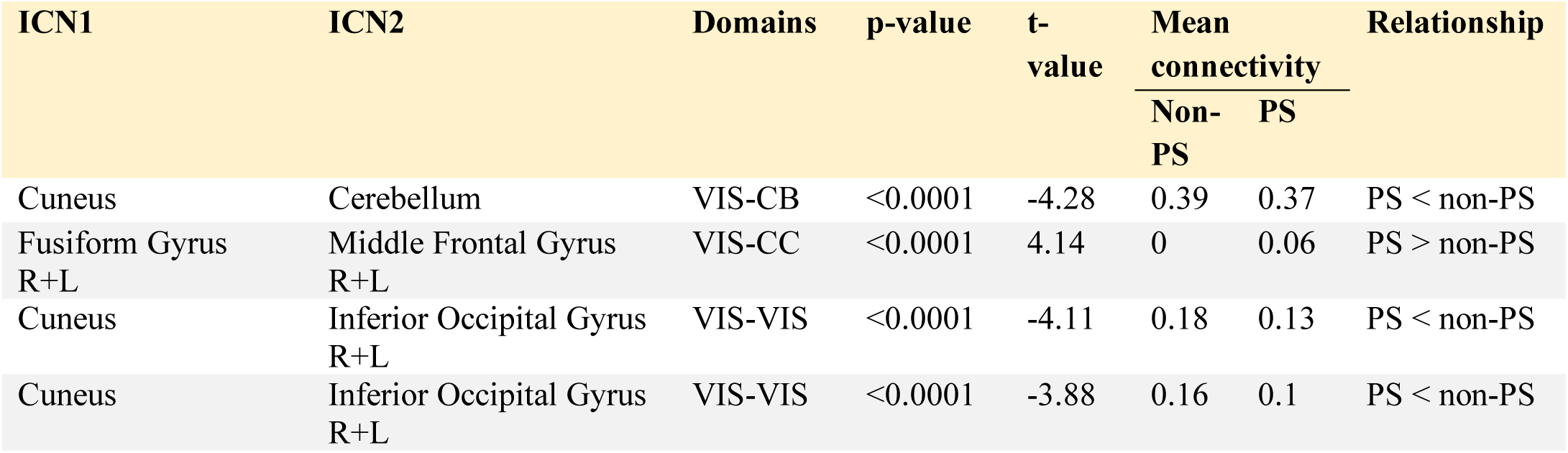
ICN-to-ICN connectivity pairs in state 5 (hypoconnected state) that show significant group effects

### Dynamic indices – Mean dwell time (MDT) and fraction of time (FT)

The factor group was not included in either model. Age was negatively associated with the time spent, overall and before transitioning to another state, in states 1 (DMN-CC domain-synchronized state) and 5 (hypoconnected state): younger participants spent more time in these states. FT and MDT increased with age in states 3 (DMN-CC domain-synchronized state) and 4 (hyperconnected state with subcortical antagonism). Results for main effects of additional covariates are presented in the Supplementary Material.

## Discussion

Here, we conducted the first analysis of whole-brain dynamic FNC in youth experiencing PS relative to their peers, revealing several novel findings. First, PS-associated altered connectivity was primarily present in states characterized by synchronization or antagonism of the DMN and CC domains. We extend previous static functional connectivity findings of hyperconnectivity between DMN, salience, and CC domains^11,12^ by showing that this pattern is state-dependent; it only occurs in a state characterized by synchronization of the DMN and CC domains, a state that also becomes less frequent with age. Further, dysconnectivity of more basic domains (sensorimotor and visual systems) is revealed in states 3, 4, and 5, completing the picture of whole-brain dysconnectivity patterns associated with PS.

### Dynamic FNC

Overall, the most notable difference between whole-brain connectivity patterns of dynamic states across groups is that state 1 is accompanied by positive connectivity between the DMN and the CC domain, whereas state 3 shows antagonism between the DMN and CC, salience, and sensorimotor domains. Changes in connectivity between the DMN and CC domains are important for adapting to cognitive demands,^52,53^ and the anterior insula, a major component of the salience domain, has been suggested to modulate connectivity between these domains.^54,55^ States 1 and 3 therefore capture snapshots of changing connectivity between the DMN and the CC domains.

By applying multivariate model selection, we found that different sets of covariates were selected for each one of the 5 dynamic states, indicating that group, sex, age, and maternal education have differential contributions to the variance of FNC across different states. The group variable was included in all but one model, and most of the differences between PS and non-PS youth occurred in states 1 and 3.

Developmental rs-fMRI studies of static FNC have shown that connectivity between the DMN and CC domains decreases with age, whereas connectivity *within* these domains increases with age;^56^ increased intra-domain connectivity of both domains was also associated with higher IQ.^56,57^ Moreover, cross-sectional studies of dynamic FNC across late childhood and adolescence also report increasing FT and MDT with age for states exhibiting DMN-CC antagonism^58^. Rashid et al.^59^ recently showed that age-related changes in ICN-to-ICN connectivity and dysconnectivity associated with autistic traits were found primarily in a state showing antagonism between the DMN and CC domains, highlighting the likely relevance of these domains to broader psychopathology.

In line with these findings, we found that MDT and FT of state 3 (DMN-CC domain-antagonized state) increased with age, whereas MDT and FT of state 1 (DMN-CC domain-synchronized state) decreased: older participants tend to spend more time in states exhibiting antagonism between DMN and CC domains and less time in states characterized by synchronization of these domains.

Collectively, these findings suggest that there may be a link between delayed neurocognitive development in PS youth^10^ and our findings of focused dysconnectivity in states generally associated with brain maturation and cognition. Future prospective studies are warranted to investigate longitudinal associations between cognitive maturation and dynamic FNC metrics, and how these factors relate to emerging PS over time.

## State 1 The DMN-CC domain-synchronized state

In state 1, PS youth exhibited hyperconnectivity of prefrontal brain areas assigned to the salience, CC, and DMN domains relative to non-PS participants. In a recent investigation of the association between static FNC and multiple dimensions of psychopathology in this cohort, PS were associated with increased connectivity between the DMN, salience, and CC domains.^12^ Further, a recent study of dynamic FNC in a CHR cohort found less temporal variability of functional connectivity in regions of the DMN and salience domains relative to healthy controls.^60^ PS-associated dysconnectivity between the DMN, salience and CC domains^55,61–64^ may be a neural underpinning of the aberrant salience theory of schizophrenia.^65,66^ Briefly, this theory posits that naturally occurring internal and external stimuli that are competing for ‘attention’ are falsely categorized as salient, ultimately leading to psychotic symptoms.^63^ The insula, a major component of the salience domain, not only detects salient stimuli but also orchestrates connectivity between the DMN and CC domain in response to those stimuli.^54,63^

We also found that, in this state, PS youth showed long-range hypoconnectivity between prefrontal CC areas and the cerebellum and within the DMN. Interestingly, prior studies in healthy adults have associated stronger prefrontal-cerebellar connectivity with better executive functioning.^67^ This is another interesting target for future studies.

Relative to non-PS youth, those with PS also exhibited hypoconnectivity in state 1 between the basal ganglia and the sensorimotor domain. Subcortical-cortical dysconnectivity has been implicated in the psychosis spectrum.^38,68,69^ Recent findings suggest an association between disruption of the cortico-basal ganglia loop and motor impairments in patients with schizophrenia.^70–72^ Moreover, behavioral data indicate that abnormal involuntary movements are linked to psychosis risk in youth,^73^ and cortical-subcortical dysconnectivity may be a contributing factor.

## State 3 DMN-CC domain-antagonized state

State 3 was the most common state in both groups. Here, dysconnectivity in PS participants primarily involved visual and sensorimotor domains. A substantial body of literature indicates alterations in visual processing in schizophrenia.^74-77^ Moreover, behavioral studies in the offspring of patients with psychosis spectrum disorders indicate an association between early visual abnormalities and later development of psychosis.^78,79^ Aberrant functional connectivity of the visual domain — which we also observed in states 4 and 5 — might underlie these early perceptual processing impairments associated with PS.

The visual domain was also mainly affected in the group by age interaction in state 4, the hyperconnected state with subcortical antagonism: Connectivity between visual association areas and the angular gyrus showed age-associated decreases in PS youth, which was not observed in non-PS youth. In contrast, connectivity between auditory association cortices and the inferior frontal gyrus increased in PS youth with age which, again, was not observed in non-PS youth. In accordance with these findings, it has been shown that multisensory integration, a function of association cortices, is disrupted in patients with schizophrenia.^80,81^

In summary, our findings of dysconnectivity in sensorimotor, visual, and association cortices could map onto the hypothesis that sensory and motor signs and multisensory integration deficits are indeed among the earliest impairments along the psychosis spectrum.^9^

### Dynamic indices

There were no main effects of group for MDT and FT, suggesting that changes in these global metrics may only be detectable at the severe end of the psychosis continuum.^35,38^ This notion is supported by previous studies; whereas patients with chronic schizophrenia spent significantly more time in hypoconnected states relative to healthy controls,^38^ CHR individuals did not differ from healthy controls in MDT/FT.^35^

## Limitations

Even though PS youth have an increased risk for developing overt psychosis^1,6^, most of them will not develop a psychotic disorder: Longitudinal data are currently not available, but will be essential to understand the development and progression of the psychosis continuum, as well as factors contributing to heterogeneity in outcome. Also, longer rs-fMRI scans may allow more stable FNC estimations.^51,82–84^ Conclusion

This study provides the first evidence that dynamic functional dysconnectivity is present even at the less severe end of the psychosis continuum, complementing previous work on help-seeking and clinically diagnosed cohorts representing the more severe end of this spectrum.

Taken together, dysconnectivity observed in state 1 highlights networks previously associated with higher-order cognitive impairment in individuals on the psychosis spectrum;^11,12^ alterations in other transient states reveal alterations in more basic domains like the visual system. Metrics of time-varying functional connectivity offer promise as future diagnostic or prognostic indicators and potential targets for therapeutic interventions.^29,38,85–89^

## Acknowledgment

We thank Ruby Tow, Alex Dib, Elizabeth Riddle, Kaitlyn Hart, Miranda Madrid, Mengtong Pan, Claire Waller, and Molly Patapoff for their help with data cleaning. Data were downloaded from the database of genotypes and phenotypes (dbGaP, phs000607.v1.p1, first PNC release, C. E. Bearden, #7147). This research was supported by National Institute of Mental Health (NIMH) grants R01 MH107250 (CEB, RAO), R01 MH101506 (KHK), K01 MH112774 (MJ), and K99 MH116115 (LMOL).

